# Disentangling reporting and disease transmission

**DOI:** 10.1101/234658

**Authors:** Eamon B. O’Dea, John M. Drake

## Abstract

Second order statistics such as the variance and autocorrelation can be useful indicators of the stability of randomly perturbed systems, in some cases providing early warning of an impending, dramatic change in the system’s dynamics. One specific application area of interest is the surveillance of infectious diseases. In the context of disease (re-)emergence, a goal could be to have an indicator that is informative of whether the system is approaching the epidemic threshold, a point beyond which a major outbreak becomes possible. Prior work in this area has provided some proof of this principle but has not analytically treated the effect of imperfect observation on the behavior of indicators. This work provides expected values for several moments of the number of reported cases, where reported cases follow a binomial or negative binomial distribution with a mean based on the number of deaths in a birth-death-immigration process over some reporting interval. The normalized second factorial moment and the decay time of the number of case reports are two indicators that are insensitive to the reporting probability. Simulation is used to show how this insensitivity could be used to distinguish a trend of increased reporting from a trend of increased transmission. The simulation study also illustrates both the high variance of estimates and the possibility of reducing the variance by avE. O’Dea eraging over an ensemble of estimates from multiple time series.

## 1 Introduction

Early warning signals (EWS) are certain statistical indicators that have shown promise of predicting catastrophic changes to complex systems (Pace et al 2017; Chen et al 2014; Scheffer et al 2015). Such predictions are challenging because changes in the equilibrium state of the system are typically very small prior to the catastrophic change in the equilibrium. A key insight that the EWS approach brings to the problem is that even when the equilibrium value of the system does not change, the rate at which deviations from the equilibrium decay can decrease a great deal and this slowing down can indicate an approaching catastrophe. In fact, such slowing down can also be expected to occur for some non-catastrophic changes to the system’s equilibrium (Kéfi et al 2013), as it has its mathematical basis in the normal forms of many types of bifurcations (Kuehn 2011). As such, a literature is growing that studies application of EWS for the transition from the disease-free to the endemic equilibrium of simple compartmental models of an infectious disease (O’Regan and Drake 2013; O’Regan et al 2015; Miller et al 2017; Brett et al 2017; Drake and Hay 2017).

This literature has yet to address two seemingly unrelated problems with such an application of EWS. The first is that the theoretical basis for EWS has been based on the assumption that data on the number of infected individuals at various time points are available, which often is not the case. Often, the available data consist of reports of some number of infected individuals that have received treatment and stopped transmitting within some time period. We refer to this type of data as case reports.

A second problem is that many EWS are based on second order properties of time series (e.g., variance and autocorrelation) which have no obvious advantage over using a simple first order indicator such as a rolling mean for the simple compartmental models studied. If the disease-free equilibrium is stable and there is occasionally a randomly imported infection from some other population, the expected number of cases per unit time is a small number that grows as the system approaches the epidemic threshold. Thus it would seem sufficient to monitor the distance to the epidemic threshold by monitoring the mean number of cases per unit time, which in this situation provides an estimate of slowing down. If the endemic equilibrium is stable, the mean number of cases per unit time does not provide an estimate of slowing down but it does provide an estimate of the equilibrium, and the equilibrium moves gradually to zero as the threshold is approached. Thus in either situation, it would seem that the mean is the natural choice of indicator.

Here we derive a theoretical expectation for the mean and second order indicators based on case report data, and we also find a new reason to use second order indicators: some observational biases that affect the mean do not affect second order indicators. In the following we establish this fact for a model of case report data that is constructed by adding an observation model to a birth-death-immigration model. We then present a simulation study that illustrates how this insight could be used to distinguish between increased reporting and increased transmission of an infectious disease.

## 2 Methods

First, we review past analytic results that are applicable to the moments of the number of case reports and present them in such a context. We then extend these results to the case of a negative binomial model of observed reports, which is a standard model used when analyzing case report data. We then describe simulations used to verify our analytic results. Finally, we describe the design of a simulation study which evaluates the idea of using moment estimates to monitor the distance to the epidemic threshold in a manner that is not susceptible to confounding by trends in the reporting probability.

### 2.1 Equations for moments of the number of cases

We model the number of infectious individuals for a disease as a birth-death-immigration (BDI) process. This modeling approach is often considered appropriate for infections that are introduced into a population from time to time but are unable to spread to a great enough extent within the population to affect population-level immunity. To be clear, the idea is to map the importation of a disease to immigration, transmission within the population to birth, and the removal of infectious individuals from the transmitting population to death. The birth and death rates are assumed to depend linearly on the population size. This BDI process captures the self-exciting nature of infectious disease dynamics and yet remains analytically tractable (Bartlett 1956).

Although analytic results for the population size of BDI processes have long been known (Kendall 1948), analytic results that describe the distribution of case reports are comparatively obscure. For notifiable infectious diseases, notification data often consists of the number of individuals that were diagnosed with the disease by healthcare workers over the course of some reporting period. Based on the assumption that healthcare workers direct the infected individuals to avoid contact with susceptible individuals (Emerson 1937), a common assumption is that the number of case reports corresponds to the number of individuals removed from the transmitting population over the course of the reporting period. Hopcraft et al (2014) derived equations for a BDI process that correspond to the moments of these removals. In the remainder of this subsection, we briefly review the derivation of these equations and provide an explicit connection between them and some proposed indicators of disease emergence.

Let *P_N,n_*(*T*) denote the probability that a total of *n* infectious individuals are removed from the population over an interval of length *T* and that there are *N* infectious individuals remaining at the end of the interval. The master equation is

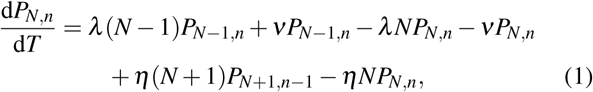

where *λ* is the per-capita transmission rate, *η* is the percapita removal rate, and *v* is the importation rate (Hopcraft et al 2014).

To obtain a solution for the moments of *n*, we make use of a moment generating function of *P_N,n_*(*T*). As we will be making use of several results from Hopcraft et al (2014), we use their definition of a moment generating function:

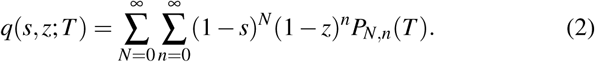

The master equation leads to a partial differential equation for the value of the generating function. For the case that *λ* < *η* and that the underlying BDI process has reached stationarity, the solution is

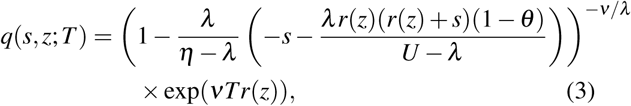

where 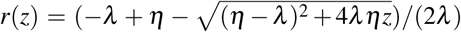, *U* = *η* − *λ*2*r*(*z*), and *θ* = exp(−(*U − λ*)*T*). By setting *s* = 0, we obtain the probability generating function for the marginal distribution of *n*. This function may be written as (Hopcraft et al 2014)

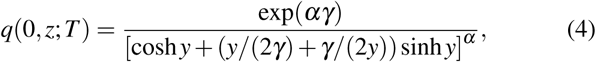

where α = *v*/*λ*, *γ* = (*η* − *λ*)*T*/2, and 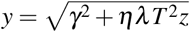. We can now calculate the mean of *n* as

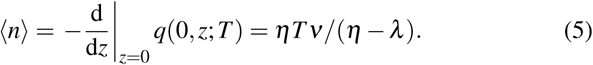

We will calculate the variance of *n* by way of the normalized second factorial moments. It turns out that the equations for the factorial moments are simpler than those for the regular moments, and later we shall see they may have some advantages as indicators of threshold distances. The normalized Rth factorial moment of a random variable *X* is defined as

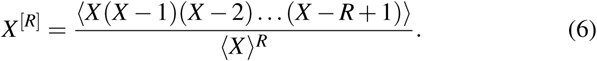

The normalized second factorial moment of *n* is then

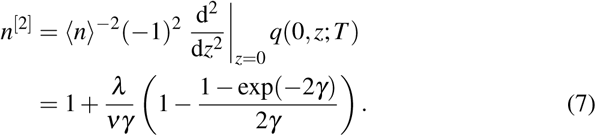

The variance follows as

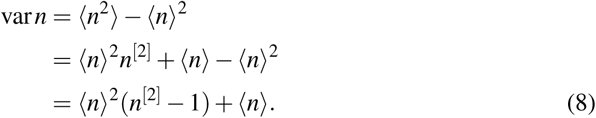

A bilinear moment function is defined by Hopcraft et al (2014) as

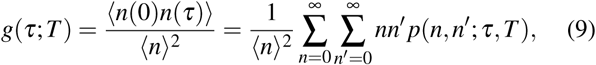

where *τ* ≥ *T* and *n*(*x*) is the number of removals that occur in the interval [*x*, *x* + *T*). The probability may be calculated as

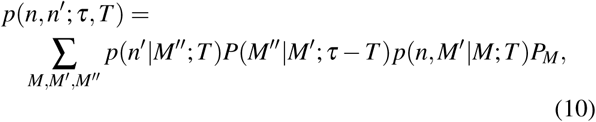

where *P_M_* is the probability of there being *M* individuals in the population at time 0 and *P*(*M*″|*M*^′^; *τ* − *T*) is the probability that the population transitions from *M*^′^ to *M*″ individuals in the time between the intervals in which *n* and *n′* are counted. In Hopcraft et al (2014), *g* is derived in terms of the moments as

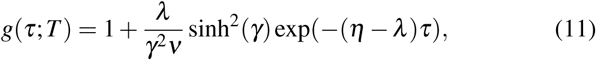

where as above we have assumed that the process is observed after reaching stationarity. A stationary autocorrelation *ρ* for a series of observations with reporting intervals of length *T* can of course be written in terms of *g* as

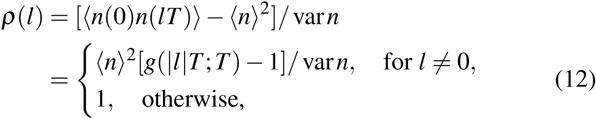

where *l* is an integer lag.

### 2.2 Binomial model for case reports

The moment equations derived in the previous subsection have been derived under the assumption that every removal is observed as a case report. Such a complete record of the process is not a realistic model for the data from most surveillance systems. It is more realistic to suppose that each removal is reported only with a given probability. That is, a step toward realism is to assume that the number of case reports *m*_bin_ is binomially distributed, with *n* being the number of trials and *ξ* being the probability of success in each trial. The binomial distribution may be a good model for the number of reported cases in which each case report corresponds to an individual who received medical care, was tested for the disease, and then prevented from having contact with others, thereby being removed from the population of transmitting individuals. For a highly specific diagnostic test, it is unlikely for the number of reported cases to ever exceed the number of true cases. The binomial preserves that upper bound on the number of reports.

For binomial sampling, the generating function for the number of case reports can be obtained by substituting *ξz* for *z* in (4), the generating function for the number of removals (Hopcraft et al 2014). With this substitution, we obtain the moments in the same manner as above, yielding

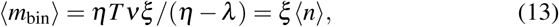

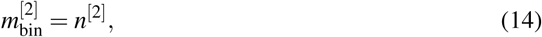

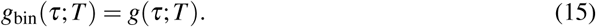

We have not provided details about the calculation of the bilinear moment because the steps of the calculation for an equivalent model (a BDI model with a rate *μ* for unobserved removals and *η* for observed removals) are outlined by Hopcraft et al (2014).

Equations (13), (14), and (15) may be used to write out equations for two standard EWS—the variance and autocorrelation:

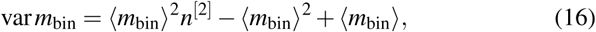

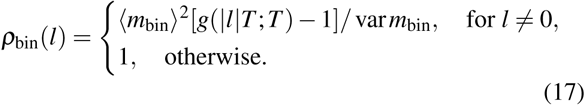

Another common EWS, the coefficient of variation, follows from (13) and (16) as

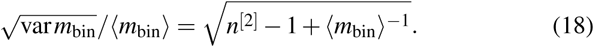

Finally, EWS are sometimes based on the power spectrum, which we denote *S*. We can obtain an equation for *S* using (17), (16), and the Wiener–Khinchin theorem:

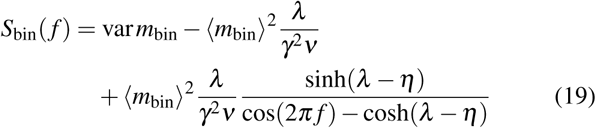

where *f* is the number of cycles of the basis function per unit time.

### 2.3 Negative binomial model for case reports

The binomial model may be an unsuitable one for the notification data of many infectious diseases. It does not allow for the number of case reports to exceed the true number of removals. In practice, over-reporting may occur due to misdiagnoses or clerical errors. For such reasons, analysts often allow for overdispersed reporting distributions when fitting case report data (Bretó et al 2009; He et al 2010; Martinez-Bakker et al 2015). Following Bretó et al (2009) we allow for such overdispersion by assuming that the case reports are negatively binomially distributed and employ the mean-dispersion parameterization of the negative binomial. The mean is assumed to be equal to the true number of case reports *n* times the reporting probability *ξ*. The variance of the number of case reports, conditional on *ξn*, is determined by a dispersion parameter *ϕ* according to *ξn* + (*ξn*)^2^/*ϕ*.

The moments of the number of case reports with this negative binomial observation model can be obtained by viewing the number of case reports *m*_nb_ as the sum of two random variables: (i) the number of removals *n* times the reporting probability *ξ* and (ii) the difference between the *ξn* and *m*_nb_, which we denote with *e* (for error term). In symbols, *m*_nb_ = *ξn* + *e*. Clearly, conditional on *n* taking a particular value, the expected value of the error term is zero. That is,

〈*e*|*n*〉 = 0. Thus the unconditional mean

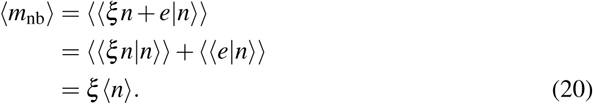

To obtain the second factorial moment, we note that 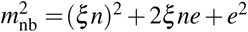. The conditional means of the terms are

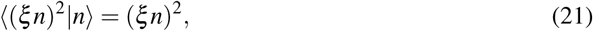

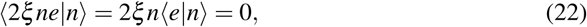

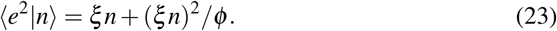

By removing the conditioning, we obtain 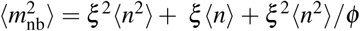. Using the above equations for the moments of *n*, we have

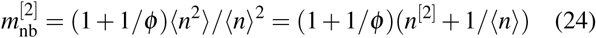

The decomposition *m*_nb_ = *ξn* + *e* also leads to an easy proof that the bilinear moment function of the negative binomial case reports must be the same as that of the removals. Consider

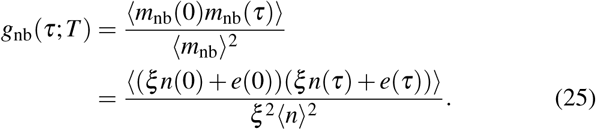

The zero conditional mean of the *e* terms and their independence from the value of *n* and *e* in adjacent reporting periods then yields

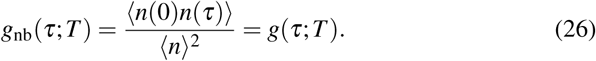

The variance, autocorrelation, coefficient of variation, and power spectrum can now be written as

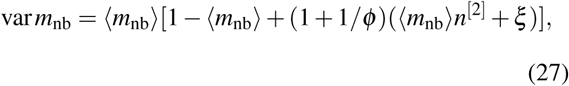

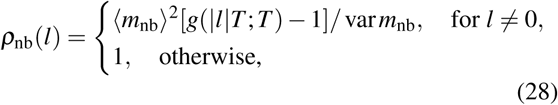

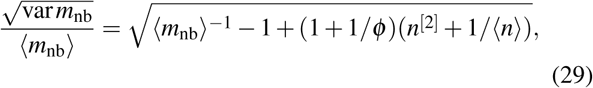

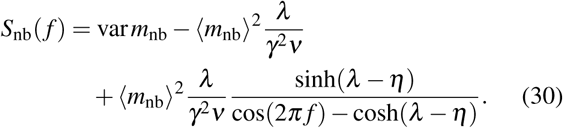

### 2.4 Numerical verification of analytic results

To verify the accuracy of the above equations for the moments, we calculated moments numerically as follows. Twenty parameter sets were obtained by uniformly sampling the removal rate *η* between 0.01 and 1, the reporting period length *T* between 1 and 10, the reporting probability *ξ* between 0 and 1, and the base-10 logarithm of *ϕ* between −1 and 2. For each parameter set, either the negative binomial or the binomial observation model was used with equal probability. To obtain a range of distances to the epidemic threshold, each parameter set was assigned a value of *λ* such that the quotient *λ*/*η* was varied sequentially from 0.05 to 0.95 in steps of 0.1 with 2 replicates of each quotient. For each parameter set, an ensemble of *N =* 10^5^ simulations of observations of a BDI process was computed. Individual simulations were initialized by sampling a population size from the stationary distribution of the BDI process and were run until the simulation time reached two reporting periods (2*T*). The essential outputs of individual simulation *i* were 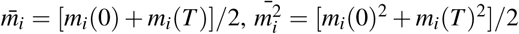, and *m_i_*(*0*)*m_i_*(*T*), where *m_i_*(0) and *m_i_*(*T*) are the number of case reports from the first and second reporting period. We numerically computed the mean as 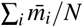, the variance as 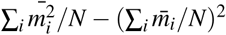, the second factorial moment as 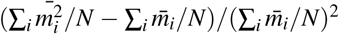, the coefficient of variation as 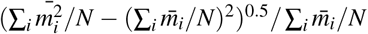, the bilinear moment function as 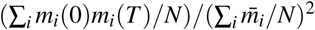, and the autocorrelation as 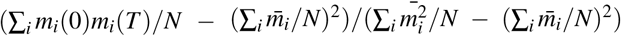. These numerical results were compared with those obtained using the above formulas by creating scatter plots.

### 2.5 Simulation study

A simulation study was performed to illustrate how these analytic results might be employed in the analysis of case reports data to distinguish trends of increased reporting from those of increased transmission. We simulated case report data over a 520 week period under four scenarios: (i) the transmission rate *λ* increases over time, (ii) the reporting probability *ξ* increases over time, (iii) *ξ* and *λ* both increase, and (iv) *ξ* decreases as *λ* increases. We then computed moving window estimates of the mean, which is expected to be sensitive to changes in both transmission and reporting, and the second factorial moment, which is expected to be sensitive to changes in transmission only. We then plotted the range of values containing the middle 90 percent of the moving window estimates as a function of time. These plots convey the feasibility of using the value of an estimate to reliably indicate a change in transmission.

Details of the simulations were as follows. Parameters shared among all scenarios were the removal rate *η* of 1 per week, the importation rate *v* of 1 per week, and the reporting period of 1 week. We first describe the scenarios with just a single changing parameter. In the increasing reporting scenario, the reporting probability *ξ* was initially 0.1 and it increased steadily to 0.5, with the linear increasing occurring throughout the course of the simulation. The transmission rate *λ* for this scenario was fixed at 0.9 per week. In the increasing transmission scenario, *ξ* was fixed at 0.5. *λ* was initially 0.5 per week and increased linearly to 0.9 per week over the course of the simulation. These parameterizations ensure that for both scenarios the expected number of case reports 〈*m*_bin_〉 was equal to 1 at the beginning of the simulation and equal to 5 at the end.

The scenarios with two changing parameters had the parameters change at the same rate as the above scenarios with a single changing parameter. In the scenario with increased reporting and transmission, the reporting probability *ξ* moved steadily from 0.1 to 0.5 and the transmission rate *λ* moved steadily from 0.5 to 0.9 In the decreasing reporting and increasing transmission scenario, the trajectory of *ξ* was reversed: *ξ* moved from 0.5 to 0.1 while *λ* moved from 0.5 to 0.9. This choice of parameters results in the expected number of cases reports 〈*m*_bin_〉 being fixed at 1 throughout the simulation.

Simulations for all scenarios were initialized by sampling from the stationary distribution of the BDI process, and all scenarios used a binomial model for case reports. We simulated 1000 replicates of each scenario.

The replicated simulations can also be thought of as a single simulation of a data set comprising case reports from many statistically identical locations. We thus refer to that set of simulations as a homogeneous ensemble. To simulate a multiple time series data set originating from a heterogeneous ensemble, we also performed a set of 1000 simulations for each scenario in which the parameters were sampled randomly for each time series in the ensemble. The sampling was uniform in bands around the parameter values for the homogeneous ensembles. Parameters *η* and *v* were sampled from (0.5, 1.5). In the increasing reporting scenario, the initial reporting probability was sampled from (0.05, 0.15) and the fixed *λ* values were sampled conditionally on *η* so that *λ/η* was uniform in (0.85, 0.95). In the increasing transmission scenario, the initial reporting probabilities were sampled from (0.45, 0.55) and *λ* was sampled so that *λ/η* was uniform in (0.45, 0.55). The same sampling scheme was used in the two additional scenarios with both reporting and transmission changing, with the location of the initial values being changed as necessary. The rate of change of either the reporting probability or the transmission rate was the same as in the homogeneous simulations. The reporting period *T* was also held at 1 week.

Moving window estimates of the mean and second factorial moment were estimated from the simulations as follows. We used a backward-looking window of 52 weeks. In each window, we calculated the sample mean and the sample second factorial moment. Ensemble estimates for a given week were generated by taking the mean of all estimates from individual windows with a final observation occurring that week. The quantiles of the individual window estimates were estimated from the empirical distribution of estimates from 1000 replicates. For the sake of computational efficiency, the quantiles of the ensemble estimates were estimated by a bootstrap method instead of simulating multiple ensemble data sets from the beginning. For estimates, we used the quantiles of a distribution of 300 ensemble estimates where each ensemble estimate was based on a bootstrap replicate of the 1000 individual window estimates.

To numerically verify the equations for the power spectrum, we also calculated the power spectrum from the Fourier transform of simulated time series with the simulation parameters matched to the beginning and end of two of the above scenarios: increasing transmission and increasing reporting. These simulations were also 520 weeks with a sampling frequency of 1 per week and replicated 1000 times. The power spectrum of individual simulations was averaged to reduce the variance of the power estimates.

### 2.6 Software and reproducibility

We conducted our simulations in *R* using the pomp package (King et al 2016). Code to reproduce our results is available at https://doi.org/10.5281/zenodo.1112362 in the Zenodo repository.

## 3 Results

### 3.1 Indicators of threshold distances not confounded by reporting probabilities

The equations for the moments derived in Methods provide insight into which moment estimates of case report data may provide an indication of the distance to the epidemic threshold. The accuracy of these equations is supported by the close agreement of simulated and calculated estimates in Figure 1. Regarding the use of these moment estimates as indicators, we propose that two key characteristics of a good indicator are, first, sensitivity to the difference between the transmission and recovery rates, *λ* − *η*, and, second, insensitivity to the reporting probability *ξ* of the reporting model. Based on this criteria, (13), which gives the mean for the binomial and negative binomial models, identifies the mean as a poor indicator. Also, the equations for the variance and coefficient of variation identify these common indicators as poor by these criteria. On the upside, (14) and (24) identify the (normalized) second factorial moment as a good indicator. These contrasts are illustrated in Figure 2.

**Fig. 1.**
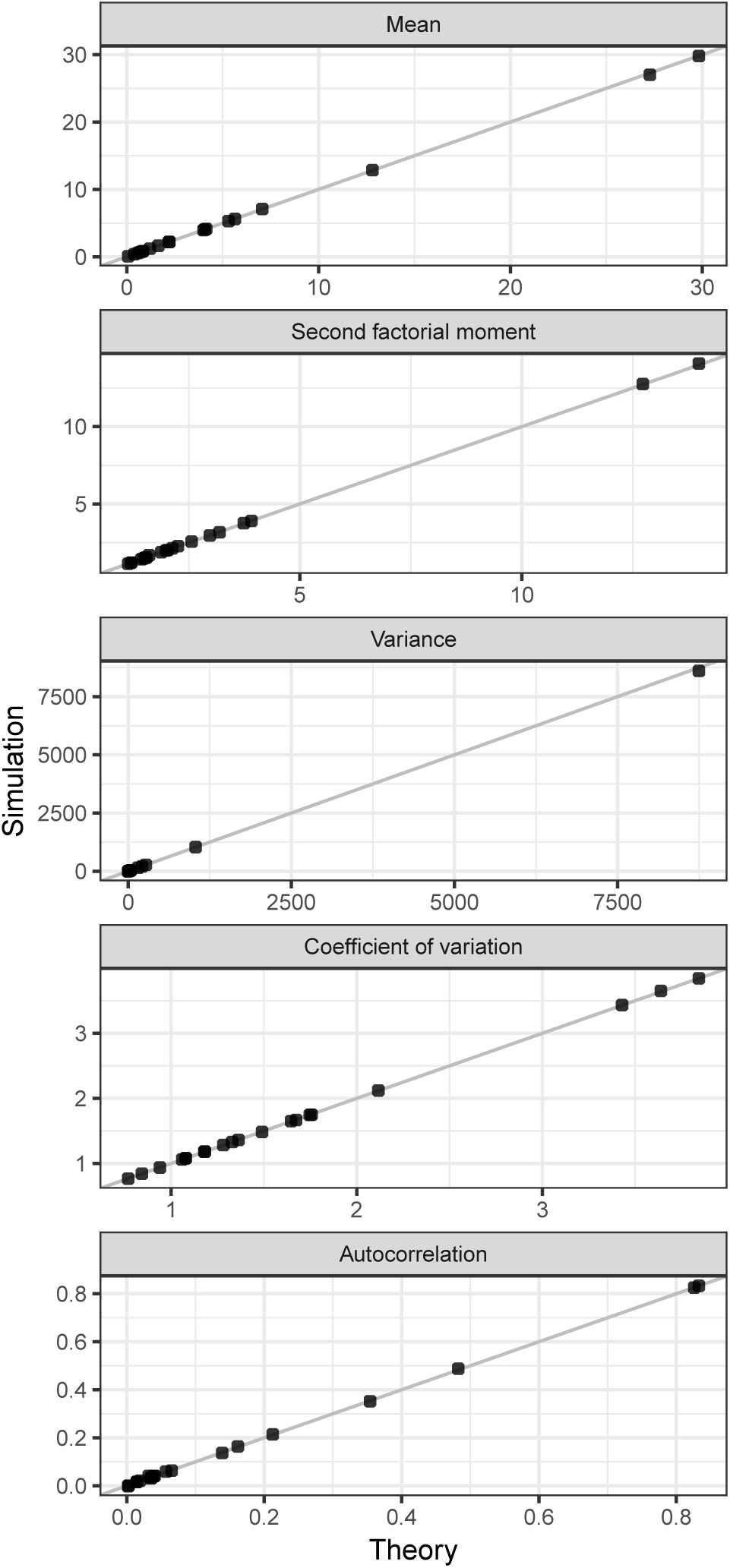
Theory and simulation agree. Model parameters were sampled as described in Methods. The line indicates where points would fall in the case of perfect agreement.

**Fig. 2.**
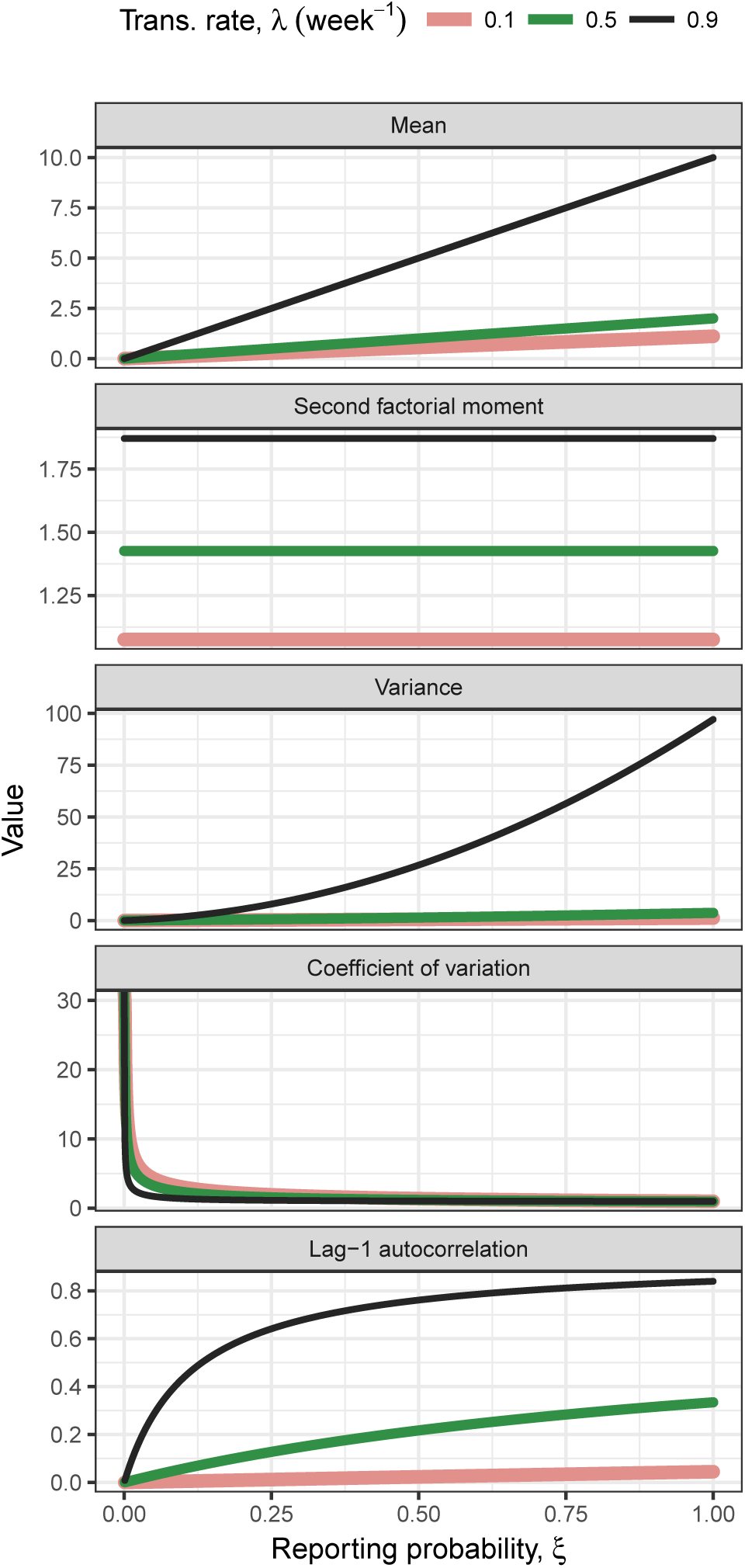
The mean is sensitive to both the transmission rate and reporting probabilities and the normalized second factorial moment is sensitive to changes in the transmission rate only. The variance, coefficient of variation, and lag-1 autocorrelation are, like the mean, sensitive to both types of changes. Lines are drawn using (13), (14), (16), (18), (17). Parameters: *η* = *ν* = 1 per week, *T* = 1 week, binomial model of reporting.

By the same criteria, one could also conclude from (11), (12), (17), and (28) that the decay rate of the autocorrelation function, which is equal to *η* − *λ*, is also a good indicator. On the other hand, the autocorrelation at a fixed lag is not a good indicator because the factor in the autocorrelation besides *g*(*τ*; *T*) − 1 is generally sensitive to *ξ*. Thus it would generally be desirable to estimate the autocorrelation function rather than a fixed-lag value of the autocorrelation function. Because this estimation is a little more complicated than estimating a factorial moment, we considered only factorial moments in our simulation study.

The case of the power spectrum is similar to that of the autocorrelation function. Equations (19) and (30) show that the power at a given frequency is generally sensitive to *ξ*. These equations also reveal that some characterization of the decay in power with frequency could be a good indicator. Figure 3 illustrates these features of the power spectrum, as well as the agreement of the equations with simulation.

**Fig. 3.**
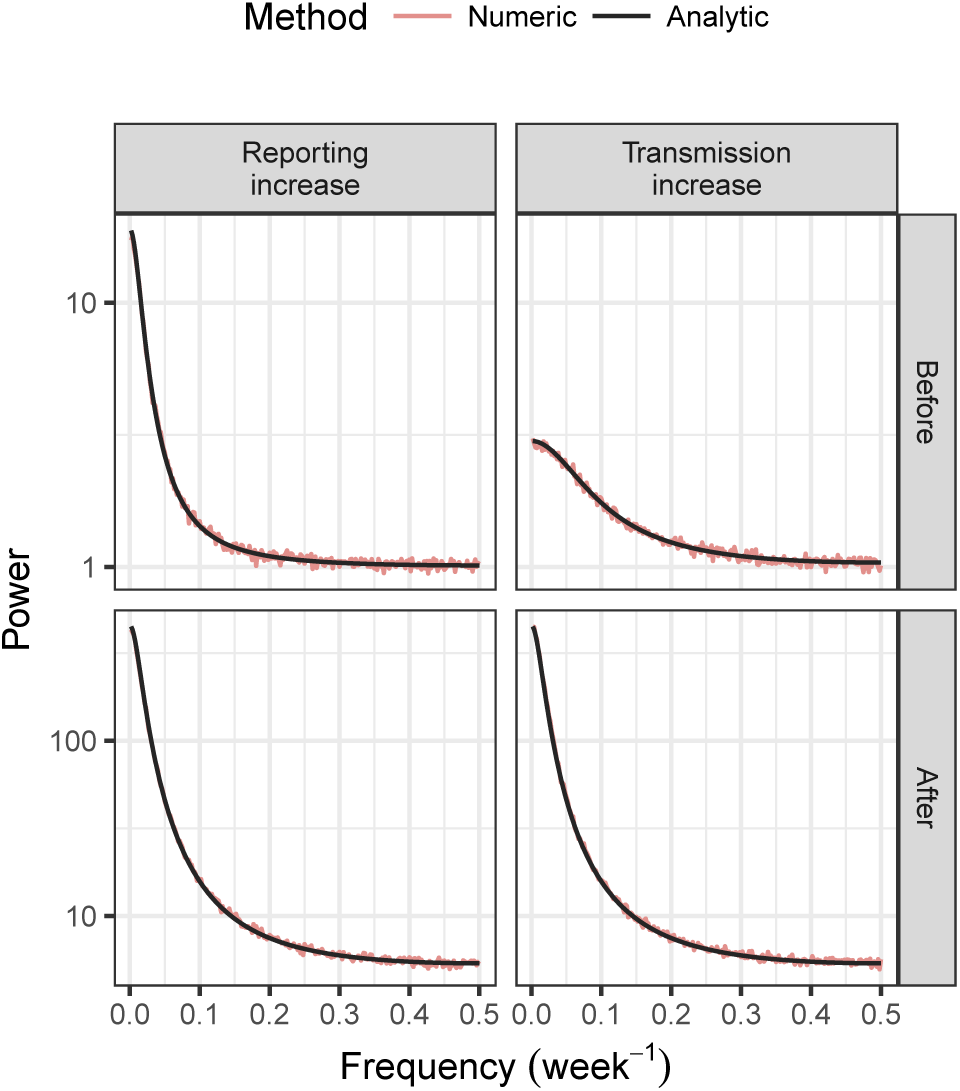
Power spectra for time series of case reports using the binomial model. Both changes to the reporting probability *ξ* and the transmission rate *λ* increase the power at a given frequency. Changes to the transmission rate affect the relative decrease in power as the frequency increases. Simulation parameters are those given for the increasing transmission and increasing reporting scenarios in Methods. The analytic power spectra were drawn using (19). The labels “Before” and “After” refer to the fixation of the parameter values to those at the beginning and end of the simulations plotted in Figure 4.

### 3.2 Application to simulated surveillance data

We next present the results of a simulation study to elaborate on the circumstances under which our criteria for good and bad indicators could prove useful. Figure 4 shows examples of the time series associated with representative simulation replicates from the scenarios describe in §2.5. Based on our analytic results, we expected that moving window estimates of the mean should increase in both scenarios, whereas increases in the second factorial moment should be specific to increases in transmission. Figure 5 confirms this expectation. However, it also conveys the extremely high variation in estimates of both moments from individual simulation replicates. For our proposed indicator of the second factorial moment, the variation of estimates around their expected values is about fivefold greater than the change in expected value due to the change in the transmission rate. Thus the variance of estimates from individual time series rules out the possibility of using this indicator to reliably signal an increase in transmission.

**Fig. 4.**
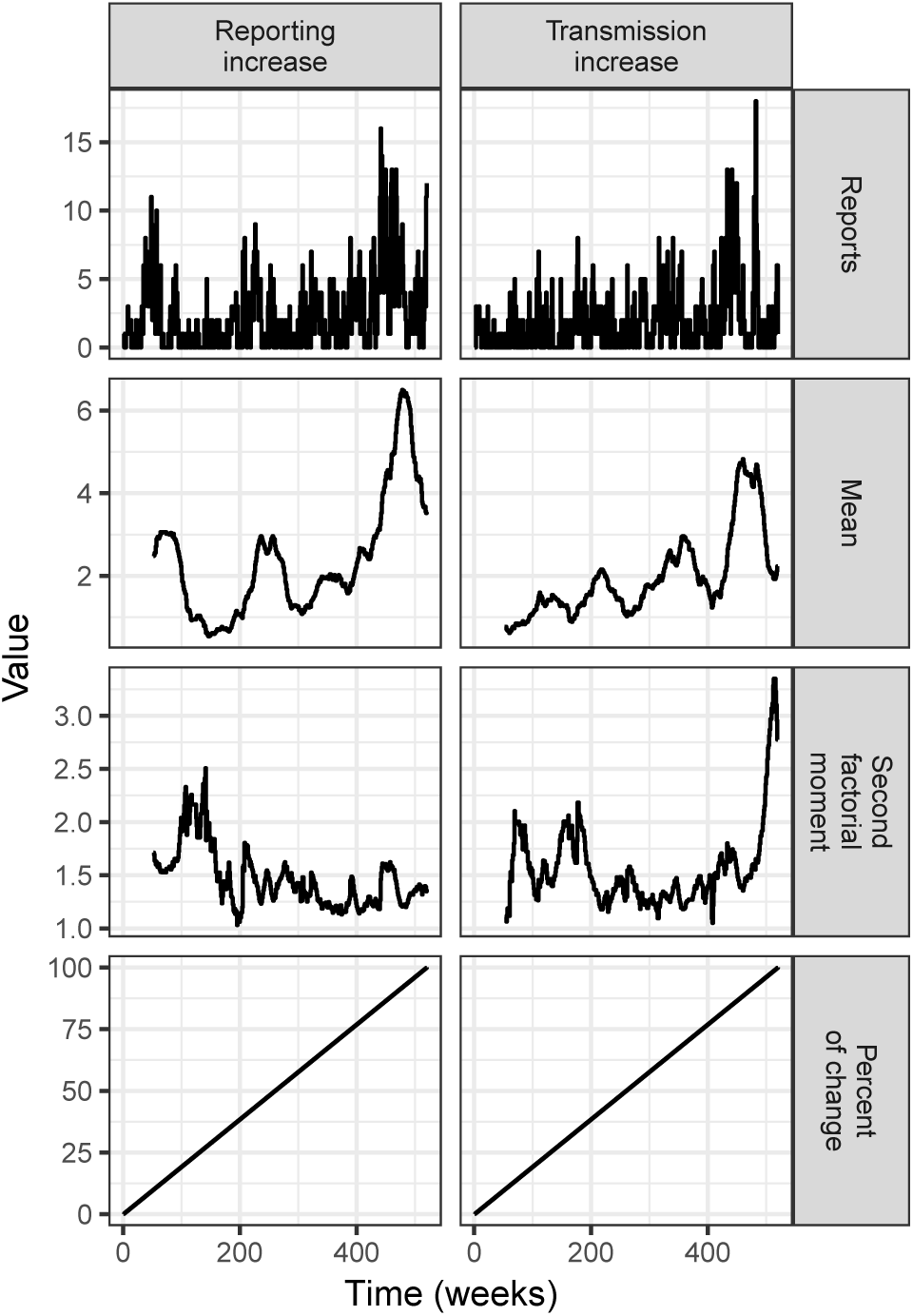
Scenarios of either increasing transmission or reporting were simulated. For one simulation of each scenario, the time series of case reports, moving window moment estimates, and the percent of the total change in the transmission or reporting parameter are plotted. Simulation parameters are those given for the homogeneous ensemble in Methods.

**Fig. 5.**
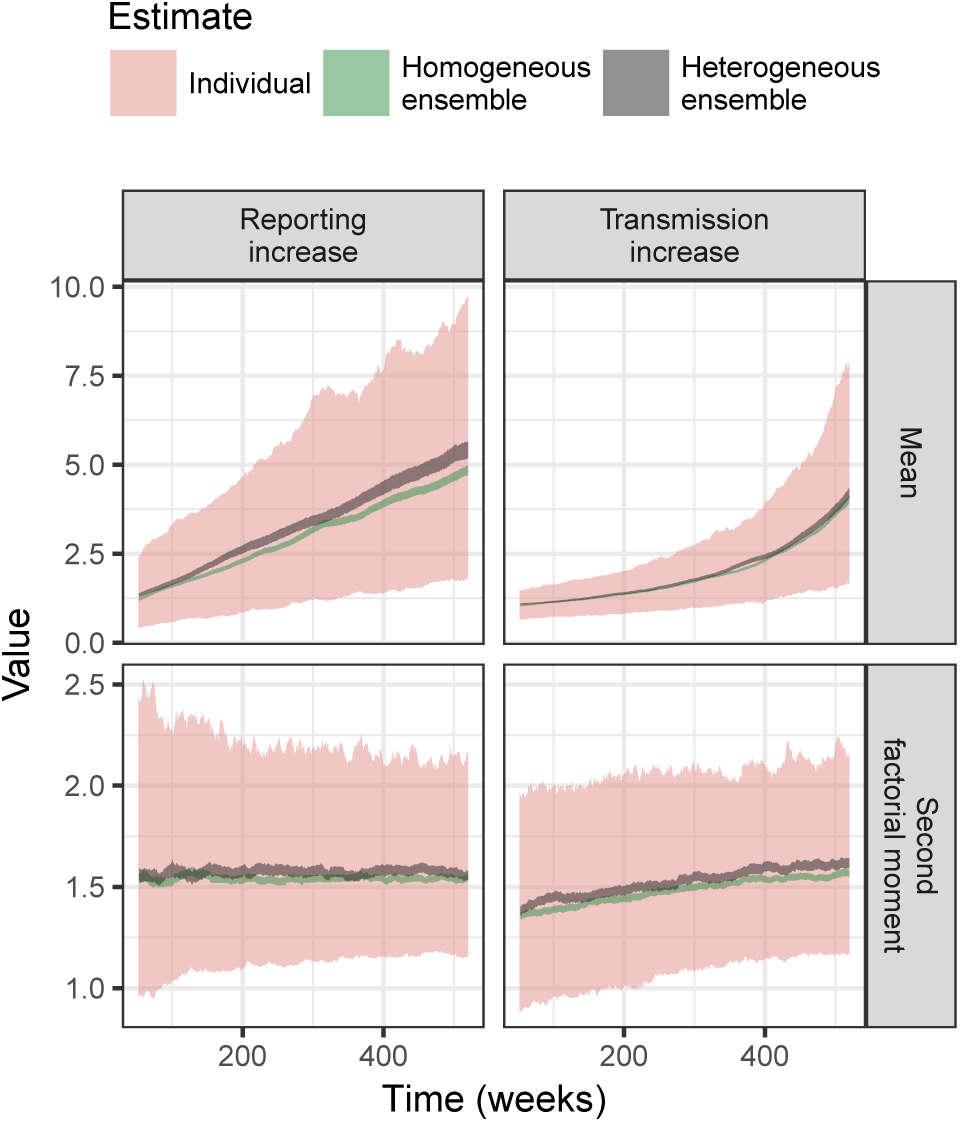
Moving window estimates of the mean increased with both reporting and transmission whereas increases in the second factorial moment were specific to increases in the transmission rate. Ribbons plot the middle 90 percent of estimates out of a set of replicated simulations. Estimates from individual time series were highly variable but ensemble estimates were much less noisy and could provide reliable information about the parameters. See Methods for the simulation parameters.

Although the application of our indicator to a single time series seems unpromising, in fact case report data are often available as multiple time series. Therefore one way to reduce the variance of the estimates would be to calculate the mean of an ensemble of moving window estimates, where each moving window estimate is based on an individual time series. For example, if one had a time series of case reports from a number of areas, one could calculate sample moments in moving windows for each area and then take the average of estimates across areas to obtain an ensemble estimate. Figure 5 shows that with an ensemble of size 1000 the signal-to-noise ratio of the second factorial moment estimates is drastically improved. A threshold of 1.5, say, on the second factorial moment could reliably discriminate between conditions of low and high transmission (weeks 1 to 250 versus weeks 250 and later).

The use of ensemble estimates raises the question of how similar members of the ensemble must be for the ensemble estimates to be useful. We do not attempt a systematic investigation of this question in this work but do confirm that perfect homogeneity is not a requirement. We simulated data for both homogeneous and heterogeneous ensembles. The parameters used to simulate individual time series in the heterogeneous ensembles were sampled from bands surrounding the set of parameters used for each time series in the homogeneous ensemble. Figure 5 shows that these heterogeneous ensemble estimates behaved similarly to homogeneous ensemble estimates.

It is easy to think of reasons why the reporting probability and transmission rate might change simultaneously. For example, as a disease becomes more common, it may become more readily identified and reported. Figure 6 shows that in this scenario, the mean is the more sensitive indicator. Naturally, however, the mean fails to provide a signal when the reporting probability drops as transmission increases. Such a scenario might occur if reduced contact tracing results in fewer cases being identified as well as less treatment and isolation of infectious individuals. In such a scenario, second order indicators are clearly superior.

**Fig. 6.**
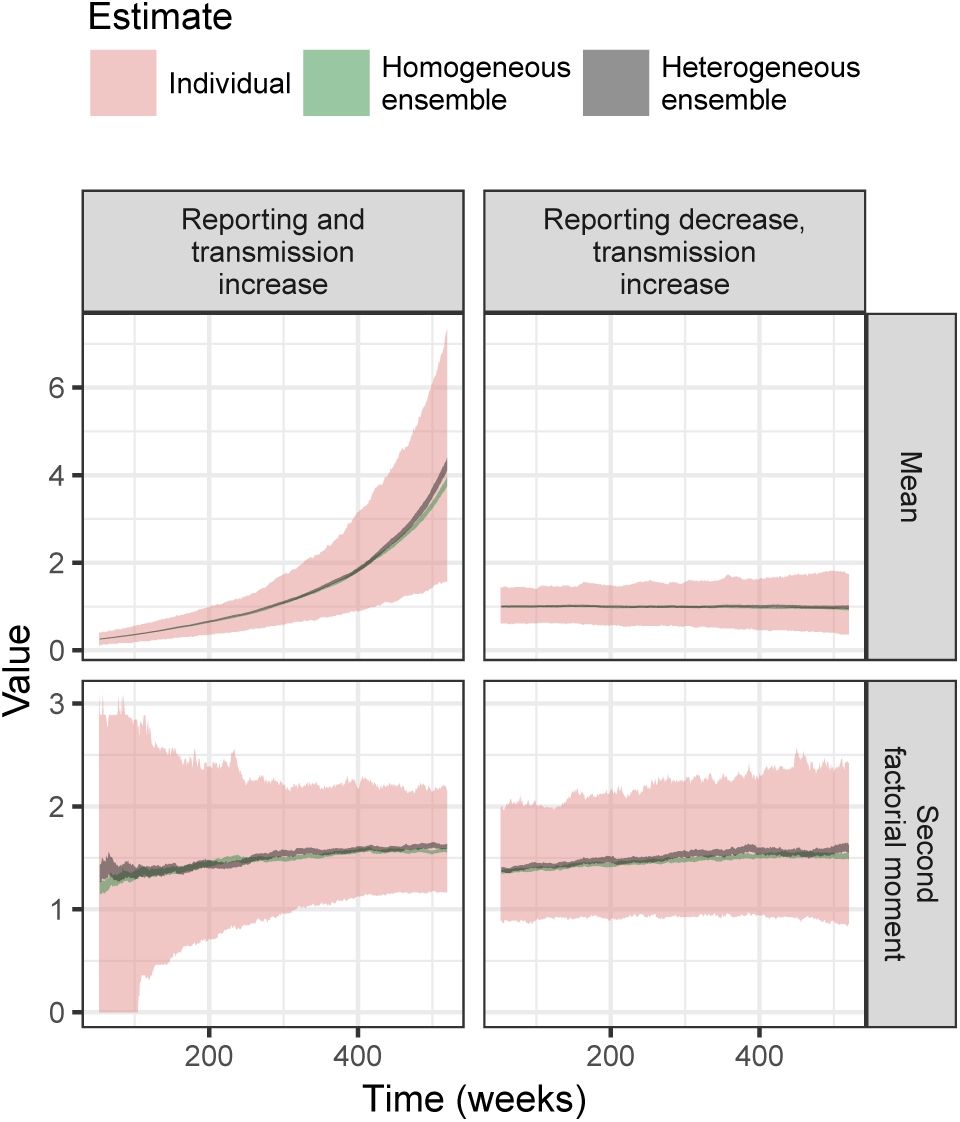
Moving window estimates of the mean can provide a more sensitive indicator than those of the second factorial moment when reporting and transmission increase simultaneously. Nevertheless, the mean fails to provide any signal when increasing transmission is counterbalanced by decreasing reporting. Ribbons plot the middle 90 percent of estimates out of a set of replicated simulations. As in figure 5 ensemble estimates had a much higher signal-to-noise ratio. See Methods for the simulation parameters.

## 4 Discussion

Our work provides analytic expressions for several moments of a model of case report data that includes mechanisms of both disease transmission and observation. These expressions make clear that certain moments are not affected by changes in the reporting probability. Our simulation results demonstrate how estimates of such a moment could be used to distinguish between trends of increased reporting and trends of increased transmission. The simulation results also illustrate that the high variance of moment estimates from individual time series presents a problem for their use as indicators. The results further show that taking the average of an ensemble of moment estimates, possibly originating from heterogeneous populations, is one possible solution. To obtain a sufficient quantity of data for such estimates, it may be fruitful to make use of digital data streams as in the works of McIver and Brownstein (2014), Richter and Dakos (2015) and Pananos et al (2017).

This work is in one respect a follow-up of the work of Brett et al (2017), which studied the behavior of the population size of a BDI (birth-death-immigration) model as it approached criticality and characterized how the slowing of the dynamics is borne out in the moments of the population size. Results for our case report models are in many ways similar to those of the BDI model. For example, the mean, variance, and autocorrelation of the case report models all increase with the approach to criticality. Our equations even imply that the decay times for the case report models, 1/(*η* − *λ*), are identical to that of the BDI model. Addition of the observation models we have considered to the BDI model, then, does not pose a fundamental problem to the idea of creating indicators to detect slowing down. It rather poses some solvable problems about trends in some indicators being confounded with trends in reporting parameters.

The problem of changes in reporting creating false impressions about trends in the transmission of an infectious disease is not specific to data analysis methods based on the slowing down phenomena. Indeed, many online aberration detection methods reviewed by Salmon et al (2016) are likely to raise alarms in response to any increase in the mean number of reported cases. Our work suggests that it may be useful to add to these systems some indicators that are based on second order moments to provide a metric of the system that may be less susceptible to changes in reporting rates. Such indicators would not require development of detailed models for each surveillance stream and would be computationally simple enough to apply at large scales.

In the literature on online forecasting of the state of outbreaks, it is well recognized that it is important to account for the random delay between when a case occurs and when it is reported (Donker et al 2011). Such a delay can lead to a decrease in the reporting probability over the most recent reporting intervals. A rapid change in the reporting probability would likely cause a change in the expected value of moving window estimates of indicators that are insensitive to the reporting probability under the assumption that the reporting probability is nearly constant within the window. However, ensemble estimates of these indicators could mitigate this problem by having small window sizes and therefore could be useful in such settings.

We caution, however, that at present any such applications should be considered highly experimental. First, it is not clear at what scale the BDI model should be considered a suitable model for the transmission of any particular infectious disease. In the BDI model we have considered, all chains of transmission share the same parameters. Although that may be reasonable at a very fine scale, case report data is typically available only at relatively large scales for which multi-type BDI models may be more appropriate. Second, it is also not clear whether our model of observation is adequate. Explicitly modeling observation and the dynamics of two frequently confused conditions, such as measles and rubella (Helfand et al 2003), may reveal that disentangling transmission from reporting is not so simple. In short, to provide a firm foundation for applications, future work must systematically determine both the conditions under which the proposed indicators perform well and whether these conditions are realistic.

Although substantial work remains to be done before it is clear what place, if any, indicators based on slowing down have to play in the routine analysis of case report data, the present work has further demonstrated the appeal of their generality. It is easy to see from our results that the decay time of the case reports will match that of the BDI process for many models of observation in which the expected value of the number of case reports is proportional to the true number of removals in the reporting period. This generality then allows us to justify decay time as an indicator for any such model. In this way, it may provide the necessary assurance that many facts that may be uncertain may not be important.

## Acknowledgements

We thank Pejman Rohani and Tobias Brett for comments on a manuscript draft. This research was funded by the National Institute of General Medical Sciences of the National Institutes of Health under Award Number U01GM110744. The content is solely the responsibility of the authors and does not necessarily reflect the official views of the National Institutes of Health.

